# Hybridization history and repetitive element content in the genome of a homoploid hybrid, *Yucca gloriosa* (Asparagaceae)

**DOI:** 10.1101/2020.06.14.150078

**Authors:** Karolina Heyduk, Edward V. McAssey, Jane Grimwood, Shengqiang Shu, Jeremy Schmutz, Michael R. McKain, Jim Leebens-Mack

## Abstract

Hybridization in plants results in phenotypic and genotypic perturbations that can have dramatic effects on hybrid physiology, ecology, and overall fitness. Hybridization can also perturb epigenetic control of transposable elements, resulting in their proliferation. Understanding the mechanisms that maintain genomic integrity after hybridization is often confounded by changes in ploidy that occur in hybrid plant species. Homoploid hybrid species, which have no change in chromosome number relative to their parents, offer an opportunity to study the genomic consequences of hybridization in the absence of change in ploidy. *Yucca gloriosa* (Asparagaceae) is a young homoploid hybrid species, resulting from a cross between *Yucca aloifolia* and *Yucca filamentosa*. Previous analyses of ~11kb of the chloroplast genome and nuclear-encoded microsatellites implicated a single *Y. aloifolia* genotype as the maternal parent of *Y. gloriosa.* Using whole genome resequencing, we assembled chloroplast genomes from multiple accessions of all three species to re-assess the hybrid origins of *Y. gloriosa.* We further used re-sequencing data to annotate transposon abundance in the three species and mRNA-seq to analyze transcription of transposons. The chloroplast phylogeny and haplotype analysis suggest multiple hybridization events contributing to the origin of *Y. gloriosa,* with both parental species acting as the maternal donor. Transposon abundance at the superfamily level was significantly different between the three species; the hybrid was frequently intermediate to the parental species in TE superfamily abundance or appeared more similar to one or the other parent. In only one case – *<u>Copia</u>* LTR transposons – did *Y. gloriosa* have a significantly higher abundance relative to either parent. Expression patterns across the three species showed little increased transcriptional activity of transposons, suggesting that either no transposon release occurred in *Y. gloriosa* upon hybridization, or that any transposons that were activated via hybridization were rapidly silenced. Further work will assess the degree to which transposon abundance and location has affected the epigenomic landscape, gene expression, and ecophysiology in *Y. gloriosa*.

## Introduction

Hybridization between related species has the potential to generate novel genotypic and phenotypic combinations, sometimes resulting in the origin of new species. Understanding the factors that promote the process of hybridization, as well as the maintenance of newly created hybrids, has been of considerable interest to both the fields of ecology and evolution (Gross and Rieseberg, 2005). As the generation of biodiversity is of primary importance to evolutionary biology, many studies have sought to determine whether or not newly created hybrids are reproductively isolated from parental species and are capable of persisting in a hybrid state for many generations. The tools aimed at studying plant hybridization include observational studies of plants and their pollinators in the wild (Leebens-Mack and Milligan, 1998; Hersch and Roy, 2007), reciprocal transplant studies across multiple environments (Wang et al., 1997), manual pollinations between related species (Sun et al., 2018), cytogenetics (Thórsson et al., 2001), and population genomics (Bredeson et al., 2016). Hybridization can result in allopolyploid individuals, in which hybridization occurs at the same time as chromosome doubling, as well as homoploids, in which there is no change in chromosome number (for a review, see Soltis and Soltis, 2009 and Rieseberg, 1997). Transposable element content and abundance has been hypothesized to contribute to genome dominance in allopolyploid species (Edger et al., 2017; Bird et al., 2018), but change in ploidy makes it difficult to assess its importance relative to hybridization in the genesis of a new species. Homoploid hybrid species provide an opportunity to focus on the effects of hybridization while controlling for ploidy level (Ungerer et al., 2009; Staton et al., 2012).

Investigation of hybridization almost always begins with a detailed understanding of the genetics and life history of the putative parental and hybrid species. In the case of wild sunflowers, numerous studies have focused on how *Helianthus annuus* and *H. petiolaris* have hybridized multiple independent times to form three homoploid hybrid species: *H. anomalus, H. deserticola,* and *H. paradoxus* (Rieseberg, 1991; Rieseberg et al., 2003). These hybrid species are morphologically distinct from their parents and each other (Rieseberg et al., 2003), display varying levels of salt tolerance (Welch and Rieseberg, 2002; Karrenberg et al., 2006), show gene expression differences (Lai et al., 2006), and exhibit population genetic patterns consistent with selective sweeps (Sapir et al., 2007). The repeated formation of homoploid hybrids in *Helianthus* has increased our understanding of hybrid speciation from both ecological and genomic perspectives, yet it is only one example of homoploid hybridization in flowering plants. Another well-studied example of homoploid hybridization is in *Iris nelsonii,* a hybrid suspected to have genetic contributions from more than two species based on patterns of both nuclear and plastid genetic variation (Arnold, 1993). The fitness of the hybrid species relatives to the parental species varies depending on the moisture of the environments, implying that genotype-by-environment interactions differentially affect parental and hybrid genotypes, a phenomenon that can lead to hybrid speciation (Johnston et al., 2001).

While hybridization’s effect on the generation of biodiversity and the movement of adaptive traits between species has been well established, the effect on the genome is only recently being fully understood. Barbara McClintock described hybridization as a “challenge” or “shock” for the genome (McClintock, 1984); the merger of two separate genomes in a single nucleus results in a completely novel genomic environment. Post hybridization, alleles once restricted to separate species now interact in a new cellular setting, allowing for the formation of novel phenotypes, epistatic interactions, and potentially significant and rapid evolutionary change. Possible outcomes of hybridization and subsequent genome shock include: alteration of gene expression (Hegarty et al., 2009; Xu et al., 2009); chromosomal rearrangements (Rieseberg et al., 1995; Lai et al., 2005; Danilova et al., 2017); genome dominance, in which one progenitor genome expresses and/or retains more genes (Rapp et al., 2009; Bardil et al., 2011; Schnable et al., 2011; Yoo et al., 2013; Edger et al., 2017; Bird et al., 2018); epigenetic perturbation (Salmon et al., 2005), which in turn can lead to a release of silencing of repetitive elements and allows for subsequent repeat proliferation (Ungerer et al., 2006; Parisod et al., 2009).

Repetitive elements in particular have been implicated in the divergence of hybrid species from their progenitors. For example, RNA-seq suggests that established homoploid hybrid sunflowers, as opposed to newly synthesized hybrids, have elevated transposon expression levels (Renaut et al., 2014). In two of these hybrid sunflower species fluorescent *in situ* hybridization studies identified expansions of *Gypsy* retrotransposons relative to the progenitor species (Staton and Ungerer, 2009). *Gypsy* and *Copia* elements are both Class I retrotransposons that replicate via a “copy and paste” mechanism (Wessler et al., 1995), in contrast to the variety of Class II DNA transposons that replicate via a “cut and paste” mechanism (Feschotte and Pritham, 2007). Transposons can affect traits by disrupting genes, duplicating or re-organizing genes (Xiao et al., 2008), or they can land upstream and create new patterns of gene expression (Studer et al., 2011). The accumulation of transposons contributes to a large proportion of genome size variation seen in plants (Tenaillon et al., 2011), and ectopic recombination between transposable elements can result in genomic deletions and are a major force in genome evolution (Devos et al., 2002).

While homoploid hybrid systems are relatively rare, recent efforts to sequence the genomes of *Yucca* (Asparagaceae) species allows us to investigate the effects of hybridization on a homoploid genome. *Yucca aloifolia* and *Yucca filamentosa* are emergent models in understanding the evolution of CAM photosynthesis, as the species use CAM and C3, respectively (Heyduk et al., 2016). The two species also hybridize to form *Y. gloriosa* (Rentsch and Leebens-Mack, 2012), which is photosynthetically intermediate and a relatively recently derived homoploid hybrid species (Trelease, 1902). All three species are sympatric in the southeastern United States, with *Y. filamentosa* found across a broader range of the eastern seaboard, including into New England and the Midwest; *Y. aloifolia* is restricted largely to the southeastern United States and reaches only as far north as North Carolina. *Yucca gloriosa* is even more restricted than either parent in its range, found only in the coastal dune systems of the Atlantic seaboard and, based on herbarium records, along the coast of the Gulf of Mexico. It is thought that *Y. aloifolia* was introduced into the southeastern United States from Mexico or the Caribbean by Spanish colonists (Trelease, 1902; Groman and Pellmyr, 2000). Perhaps as a result of the human-involved introduction, *Y. aloifolia* has escaped the dependence on the obligate *Yucca*-yucca moth pollination mutualism and can be pollinated by the yucca moth *Tegeticula yuccasella* (Leebens-Mack and Pellmyr, 2004) or introduced generalist honeybees (*Apis mellifera*) (Rentsch and Leebens-Mack, 2014). *Yucca filamentosa* still retains its obligate pollination mutualism with the yucca moths *(Tegeticula yuccasella and T. cassandra)* (Pellmyr, 1999), and overlaps in flowering time with *Y. aloifolia* briefly and only in some years, suggesting that hybridization between the two species may be rare.

Previous work suggested no variation in chloroplast or microsatellite repeats in a small sampling of *Y. aloifolia* genotypes, and further indicated that *Y. aloifolia* is the maternal parent in any hybridization events that led to *Y. gloriosa* (Rentsch and Leebens-Mack, 2012). Through a whole genome sequencing project that aims to assemble the genomes of *Y. aloifolia* and *Y. filamentosa,* resequencing was performed on a number of individuals of all three *Yucca* species.

Using the resequencing data, we sought to re-test hypotheses on the number and direction of hybridization events in *Y. gloriosa.* We further examined the repeat landscape of all three species to determine if repeat content in the hybrid is purely additive, or if transgressive repeat phenotypes exist that suggest some degree of genomic shock post hybridization. Finally, using existing RNA-sequencing datasets in the three species of *Yucca,* we examined the activity of repeats using mRNA reads as a proxy. Through the use of high throughput genomic data, we find that *Y. gloriosa* is the result of repeated and bi-directional hybridization events that evidently led to minimal repeat proliferation. Our findings further suggest that there is little evidence of repetitive element release in *Y. gloriosa* as a result of hybridization.

## Materials and Methods

### DNA sampling, library preparation, and sequencing

Clones of 41 individuals (5 from *Y. aloifolia,* 24 from *Y. gloriosa,* and 12 form *Y. filamentosa*) were collected throughout the Southeastern United States from 2013 to 2015 and planted in the University of Georgia greenhouse (Figure 1, Supplemental Table 1). In 2018, approximately 100 mg of fresh tissue was harvested from fully expanded leaves and kept on ice until DNA extraction, using a CTAB protocol with sorbitol addition that removes secondary compounds before DNA purification (Doyle, 1987; Štorchová et al., 2000). DNA was visualized on a 1.5% agarose gel to measure integrity and quantified via Qubit. Samples were shipped to the HudsonAlpha Institute for Biotechnology, where Illumina 350 basepair PCRfree fragment libraries were constructed using standard protocols. Each library was uniquely barcoded and sequenced on a NovaSeq 6000 with paired end 150bp reads. Data is available on the NCBI Sequence Read Archive (for a full list of SRA accessions, see Supplemental Table 1).

**Figure 1.**
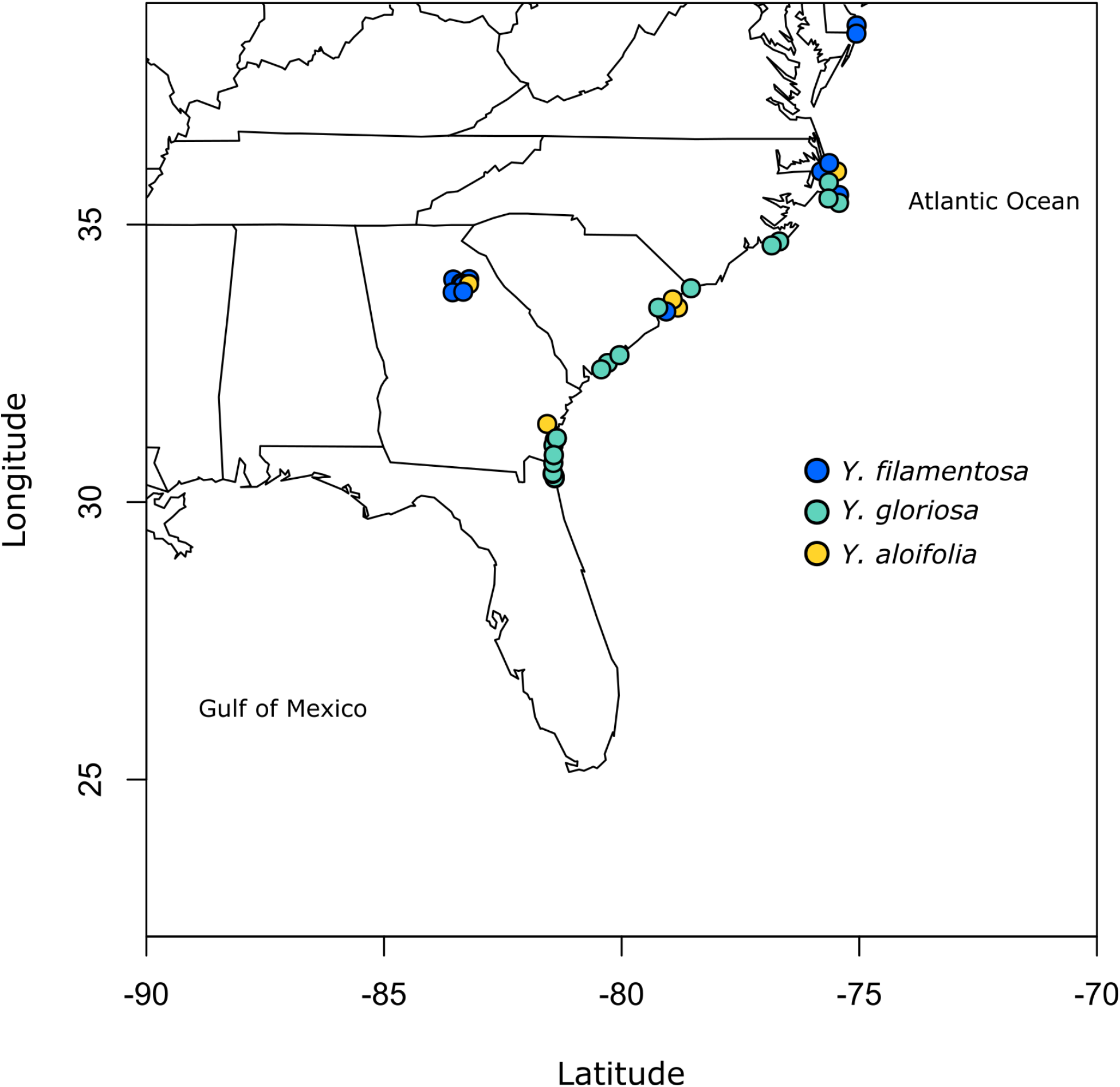
Map of populations sampled for this study. See Supplemental Table 1 for full geographic locality information.

### Chloroplast genome assembly and analysis

Raw reads were first quality trimmed using Trimmomatic v 0.36 (Bolger et al., 2014). Due to the sheer size of the sequence data per individual – roughly 400-800 million reads — a subset of four million paired-end reads was randomly sub-sampled from each library’s trimmed dataset in order to speed up computational analyses. The sub-sampled data were used as input into the program Fast-Plast (https://github.com/mrmckain/Fast-Plast), which assembles plastid genomes by first mapping reads to a reference plastid genome (here we used a previously assembled *Y. filamentosa* chloroplast genome (McKain et al., 2016)).

Chloroplast genomes of *Agave americana* (NCBI accession: KX519714.1, Abraham et al., 2016) and *Nolina atopocarpa* (NCBI accession: NC_032708.1) were used as outgroups for phylogenetic analyses. All *Yucca* chloroplast assemblies as well as *Agave* and *Nolina* were aligned using MAFFT (Katoh and Standley, 2013). The alignment was manually inspected for misaligned regions, and as a result three *Yucca* genotypes (*Y. aloifolia* YA7, and *Y. gloriosa* YG13 and YG61) containing considerable misalignments indicative of a sub-optimal genome assembly were not included in further analyses. The second inverted repeat (IR) region was removed before tree estimation: an aligned *Y. aloifolia* chloroplast genome sequence (YA23) was annotated for the IR by conducting a BLASTn (Altschul et al., 1990) against itself. The position of an inverted self-hit in YA23 was used to remove the second IR from the multi-species alignment. The optimal model of molecular evolution (GTR+Gamma) was determined using JModelTest v2 and BIC penalized-likelihood (Darriba et al., 2012) on the CIPRES gateway (Miller et al., 2010). The multiple sequence alignment was then used to estimate a chloroplast phylogeny using RAxML v8.2.11, with 500 bootstrap replicates (Stamatakis, 2006). The entire chloroplast alignment (with both IR) of the *Yucca* species without outgroups was also used to construct a median joining chloroplast haplotype network using PopArt (epsilon = 0) (Leigh and Bryant, 2015). Chloroplast genome assemblies were annotated in Geneious Prime 2019.2.3, using the built-in annotation tool with the previously published *Y. filamentosa* annotation as a reference. Chloroplast genome assemblies have been uploaded to NCBI’s GenBank, and the plastid alignment and newick files can be found at https://github.com/kheyduk/Yucca_plastome.

### Repetitive content annotation and analysis

In a similar fashion to the chloroplast sequence processing, one million trimmed paired-end reads were randomly sub-sampled for an analysis of transposon content. In order to ensure that only nuclear repetitive sequences were being analyzed, reads were first mapped to *Yucca* chloroplast and mitochondrial genome sequences (reference files are available at JGI Genome Portal, genome.jgi.doe.gov) using Bowtie v2 with default settings (Langmead and Salzberg, 2012) to be flagged for removal. The nuclear data were retained and further processed in preparation for downstream steps, including: converting bam mapping files to fastq files using SAMTools v1.9 (Li et al., 2009) and BEDTools v2.26 (Quinlan and Hall, 2010), interleaving fastq files so that pairs are found sequentially in a single file (script available at https://github.com/sebhtml/ray/blob/master/scripts/interleave-fastq.py, from Boisvert et al., 2010), and converting fastq files to fasta files with the FASTX-Toolkit v 0.14 (http://hannonlab.cshl.edu/fastx_toolkit/).

Transposome (Staton and Burke, 2015) was used to cluster and identify repetitive DNA sequences in all 41 *Yucca* genotypes using a *Yucca-*specific reference. Briefly, RepeatModeler was used to predict repeat families *de novo* on the assembled *Yucca* genomes; RepeatModeler uses both RECON (Bao and Eddy, 2002) and RepeatScout (Price et al., 2005) to identify repeat family consensus sequences. To remove false positives (e.g., repetitive domains within genes), the predicted RepeatModeler consensus sequences were searched for functional PFAM and Panther domains. If no domains – or only known transposable element domains – were found in a given putative repeat family, it was retained as a true repeat; if only false positive domains were identified, the family was removed from further analysis. Putative repeat families that had a combination of transposable element and false positive domains, or had otherwise unknown domain classes, underwent manual curation.

For annotating *Y. aloifolia* and *Y. filamentosa* repeats via Transposome, we used the species-specific RepeatModeler families (repetitive element reference files are available at JGI Genome Portal, genome.jgi.doe.gov). For *Y. gloriosa* hybrid individuals, we concatenated the two parental repeat databases. Finally, we used the following parameters in our usage of Transposome: percent identity = 90%, a required fraction of overlap between pairwise matches of 0.55, a minimum cluster size of 100, a merge threshold of 1000, and a BLAST e-value of 1. Cross-species comparisons of transposon annotation included the average amount of total repetitive DNA as well as the relative amounts of the annotated transposon families. In R v. 3.6.1 (R Core Team, 2019), we used ANOVA to determine whether there were significant differences between species in the relative amount of repetitive DNA in each of the 10 annotated families. Additionally, a data matrix containing each individual’s relative amount of repetitive DNA for each of the 10 annotated families served as the input for a principal components analysis, using the prcomp() function in R.

### Repetitive element activity via mRNAseq

Many repetitive elements contain sequences that are involved in their replication and therefore are translated into mRNA; transcripts produced from these repeats can be detected by mRNA sequencing (Hollister et al., 2011; Dion-Côté et al., 2014). While read counts from mRNA sequencing are a proxy for transcription of a repeat, no assumptions can be made as to the successful integration of a repeat copy into the genome post transcription; a variety of genomic mechanisms exist to silence and degrade repetitive element-derived transcripts (Lisch, 2009; Fultz et al., 2015). Nevertheless, as a first approximation of repeat activity, we used previously published mRNA-seq data on the three species of *Yucca* analyzed here (Heyduk et al., 2019). Briefly, RNA was collected from all three species of *Yucca* growing in growth chambers set to 30 °C/18 °C day/night temperatures, with ~ 400 μmol m^-2^ s^-1^ of light at leaf level, and 40% humidity in a 12 hour day/night light regime. While the previous study further assessed gene expression under drought, here only libraries from well-watered plants taken during the daytime were analyzed. The original study used 2-3 genotypes per species, each of which had 2-3 replicates that were taken from different time points during the day. Because replication within a genotype is confounded with time, we limited our analyses to considering only species-specific differences rather than examining genotypic differences within species. Final species-level replication varied from 6 in *Y. aloifolia* to 9 in *Y. gloriosa* and *Y. filamentosa*.

RNA reads were mapped to the same repeat databases used in Transposome; *Y. aloifolia* and *Y. filamentosa* reads were mapped to each species’ specific repeat reference, while *Y. gloriosa* reads were mapped to a merged parental reference. RNA reads were mapped via Kallisto v 0.43 using default parameters (Bray et al., 2016). For *Y. gloriosa,* counts were summed in cases where both parental species had a consensus sequence for a given repeat family. Libraries were first normalized by the Trimmed Mean of M-values (TMM) (Robinson and Oshlack, 2010) as implemented in EdgeR (Robinson et al., 2010), then scaled by overall abundance of that repeat family as estimated by Transposome. To scale, a matrix consisting of all repeat abundances across all genotypes from the three *Yucca* species was scaled by the maximum abundance of all families identified by Transposome. These scaled abundance values were then used as a multiplier of the TMM normalized read counts. By normalizing by genomic abundance, expression of repeats could then be compared across genotypes and species that have varying genomic fraction of the repeat families. Once normalized and scaled, we tested for significant expression within species using a glm intercept model in the glm.nb() function in the R package MASS (Venables and Ripley, 2013), which employs a negative binomial model appropriate for count data that exhibits a degree of overdispersion. Differentially expressed repeats between species were also tested with a negative binomial model, and *post hoc* tests were done using the emmeans() function from the R package emmeans.

## Results

### Plastid phylogenetic and haplotype analyses

Despite the relatedness between the three *Yucca* species studied here, there was enough divergence between the species’ chloroplast genomes to identify highly supported clades of chloroplast haplotypes (Fig. 2). *Y. gloriosa* genotypes were found nested within three separate clades (Fig. 2). A single *Y. gloriosa* genotype, YG16, was within a clade that otherwise contained all of the *Y. filamentosa* individuals that were analyzed. Three *Y. gloriosa* genotypes (YG12, YG55, and YG56) were placed in a clade with two *Y. aloifolia* genotypes (YA23 and YA11). The remaining 18 *Y. gloriosa* genotypes were grouped with the remaining two *Y. aloifolia* individuals (YA3 and YA32).

**Figure 2.**
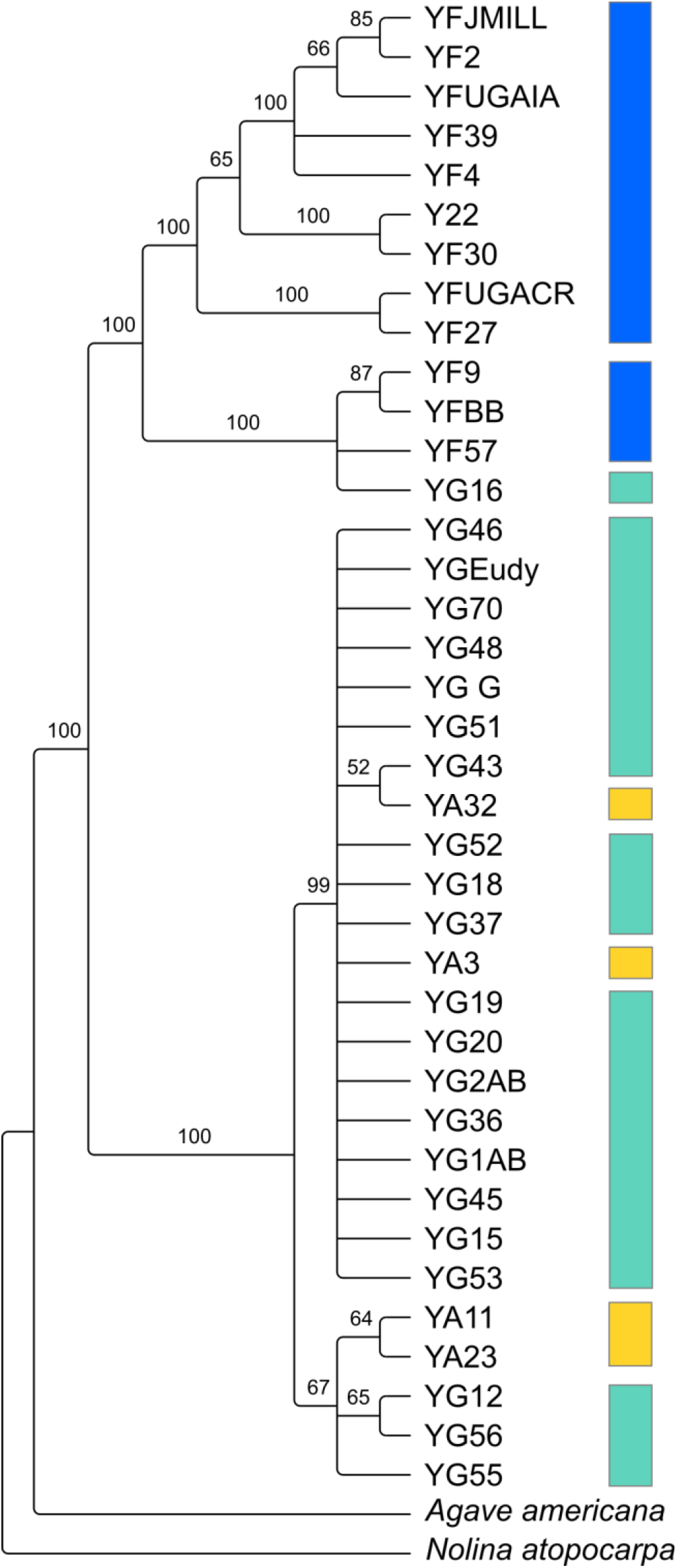
RAxML estimated phylogeny of the plastome using only one inverted repeat copy in the alignment. Bootstrap support indicated on the branches, with nodes that had less than 50 bootstrap support collapsed. Colors indicate the three species: *Y. aloifolia* (yellow), *Y. filamentosa* (blue), and *Y. gloriosa* (teal).

PopArt haplotype analysis (Leigh and Bryant, 2015) identified the same patterns found in the maximum likelihood-based phylogeny. Over 350 substitutions differentiated the two major groupings of genotypes (*Y. aloifolia* and *Y. filamentosa-like* chloroplast genomes; Fig. 3). *Yucca filamentosa* had considerably more chloroplast haplotypes compared to *Y. aloifolia* (7 vs. 2, respectively; Fig. 3). In contrast to previous analysis of nuclear simple repeats (Rentsch and Leebens-Mack, 2012), genetic diversity was seen not only in the *Y. aloifolia* chloroplast genomes but also for *Y. gloriosa*, which had four substitutions separating the different *Y. aloifolia*-like haplotypes, and over 400 substitutions separating the single *Y. filamentosa*-like haplotype from other individuals of *Y. gloriosa*.

**Figure 3.**
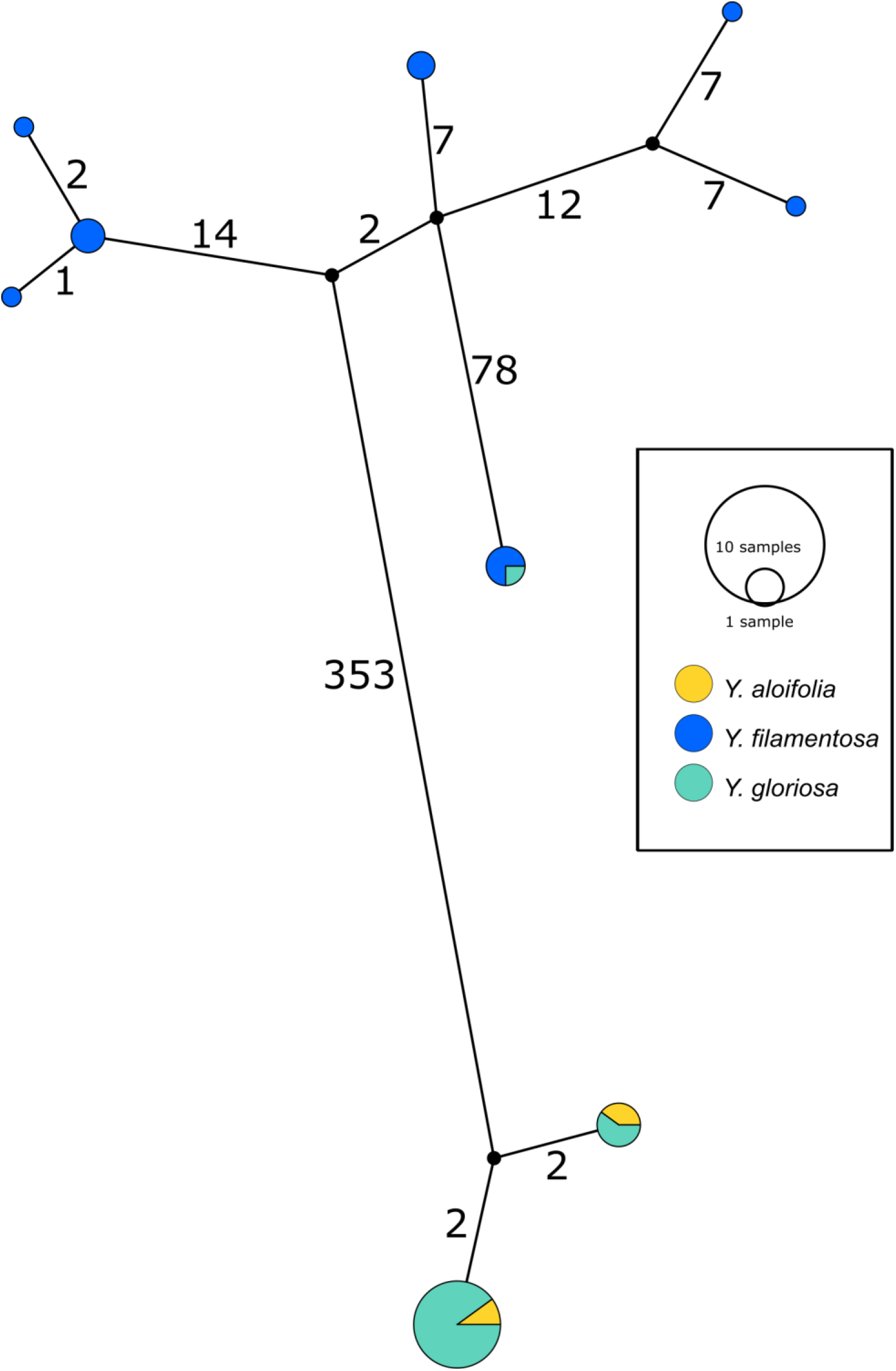
Haplotype network estimated from the entire plastome alignment across all three species, excluding outgroup accessions. Haplotype estimated via PopArt, with number of substitutions separating haplotypes on branches and size of the haplotype circles representative of the total number of individuals within that haplotype.

### *Repetitive fraction of* Yucca *genomes*

The fraction of the genome containing repetitive DNA significantly differed between the three species (p < 0.001, F2,38 = 17.853). While *Y. aloifolia* (mean repetitive genome fraction = 0.658; s.d. = 0.0138) and *Y. gloriosa* (mean = 0.662; s.d. = 0.0215) had statistically indistinguishable amount of repetitive DNA, *Y. filamentosa* was significantly lower than both species (mean = 0.621; s.d. = 0.0167; p<0.01 for both *post hoc* comparisons). Moreover, the fraction of the genome comprised of various repeat families varied across the three species. The most abundant type of repeat in all three genomes were members of the *Gypsy* superfamily (Fig. 4), comprising ~39 % of the total genome, although species did not significantly differ in overall *Gypsy* abundance. The second most abundant superfamily in the *Yucca* genomes, at about ~ 16.5 %, was *Copia* (Fig. 4). *Yucca gloriosa* had significantly more *Copia* elements than either parent *(post-hoc* comparison of *Y. gloriosa* to either parent p < 0.001). The third most abundant repeat superfamily was DNA *Helitrons,* at ~ 3.5 %, which had significantly different abundances between all three species *(post-hoc* comparison p<0.01). In general, the variation in repeat family abundance between the three species was large enough to distinguish each species (Supplemental Figure 1), though intraspecific variation in repeat abundance was apparent as well. The three *Yucca* species also exhibited presence/absence variation for repeat families: the LTR DIRS element and the non-LTR *Zisupton* elements were found in *Y. filamentosa* and *Y. gloriosa*, but not in *Y. aloifolia* (Supplemental Table 2, Fig. 4). In contrast, the LINE-2 (L2) non-LTR element, and the Novosib and P DNA elements were found in both *Y. aloifolia* and *Y. gloriosa*, but not *Y. filamentosa* (Supplemental Table 2, Fig. 4).

**Figure 4.**
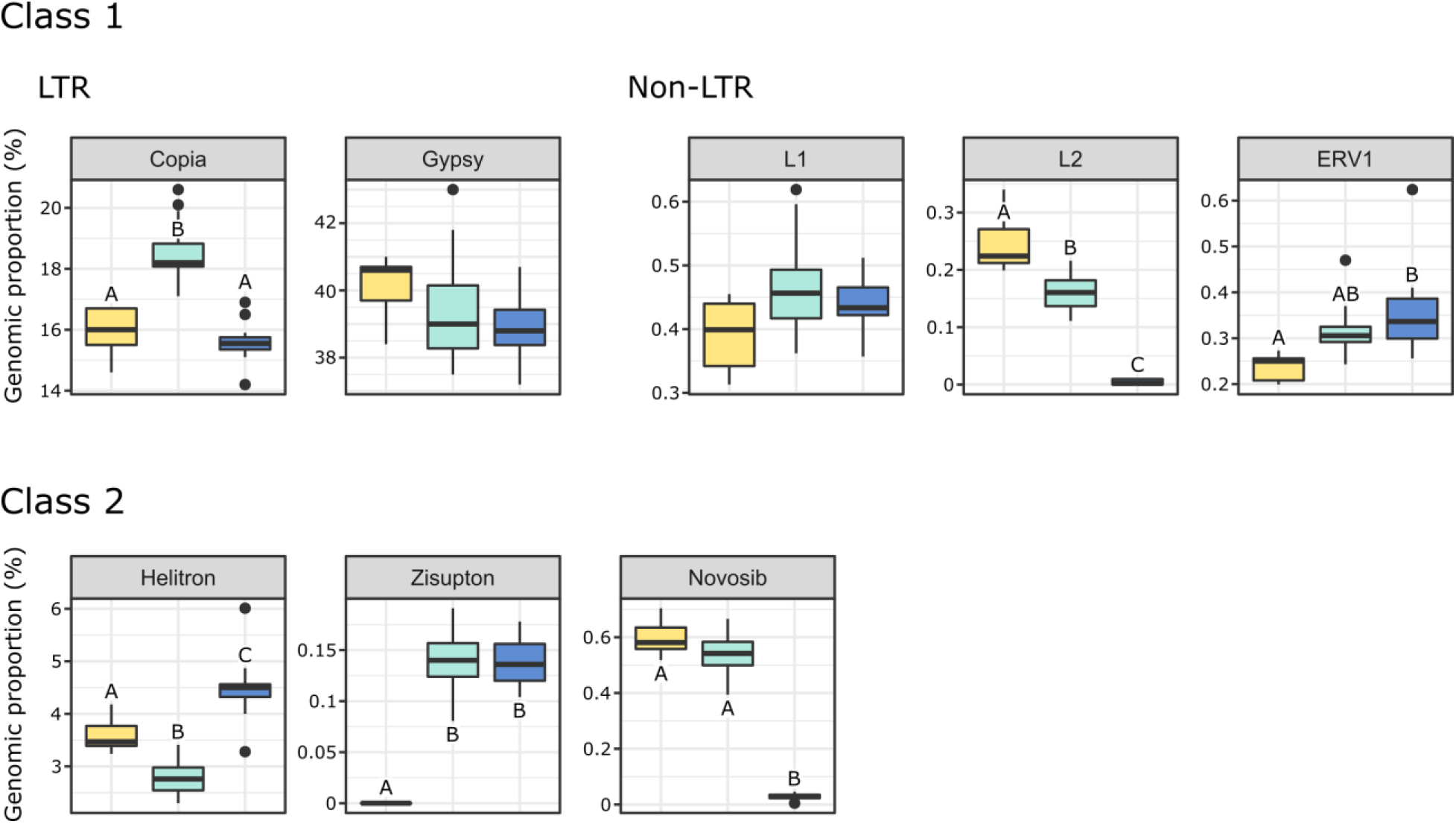
The genomic proportion (as a %) of the genome of a subset of transposable element families in *Y. aloifolia* (yellow), *Y. gloriosa* (teal), and *Y. filamentosa* (blue). Letters indicate significant differences based on Tukey *post-hoc* tests from an ANOVA (abundance ~ species) per repeat family; shared letters indicate no significant difference at p<0.01.

### Repeat mRNA expression

Transposome abundance analysis of *Y. aloifolia* and *Y. filamentosa* identified 443 and 569 repeat families in at least one genotype of either species, respectively; only 138 repeat families were present in both parental species (Fig. 5). Of the 138 families present in both species, only 118 and 119 repeat families had significantly non-zero expression in *Y. aloifolia* and *Y. filamentosa,* respectively (Benjamini-Hochberg adjusted p-value < 0.01) (Supplemental Table 3). Only 27 families were significantly expressed in both parental species (Table 1). Repeat families with significant expression were typically from *Gypsy* (64% and 61% of total families expressed in *Y. aloifolia* and *Y. filamentosa*, respectively) and *Copia* (25%, 27%) superfamilies. *Yucca gloriosa* had largely overlapping expression with its parental species; the hybrid shared significant expression of 74 families with *Y. aloifolia* and 70 families with *Y. filamentosa*. *Yucca gloriosa* had only two families that were not also significantly expressed in either parent: one a member of the *Gypsy* superfamily, the other belonging to the *Copia* superfamily, and both had genomic abundance at less than 1%.

**Figure 5.**
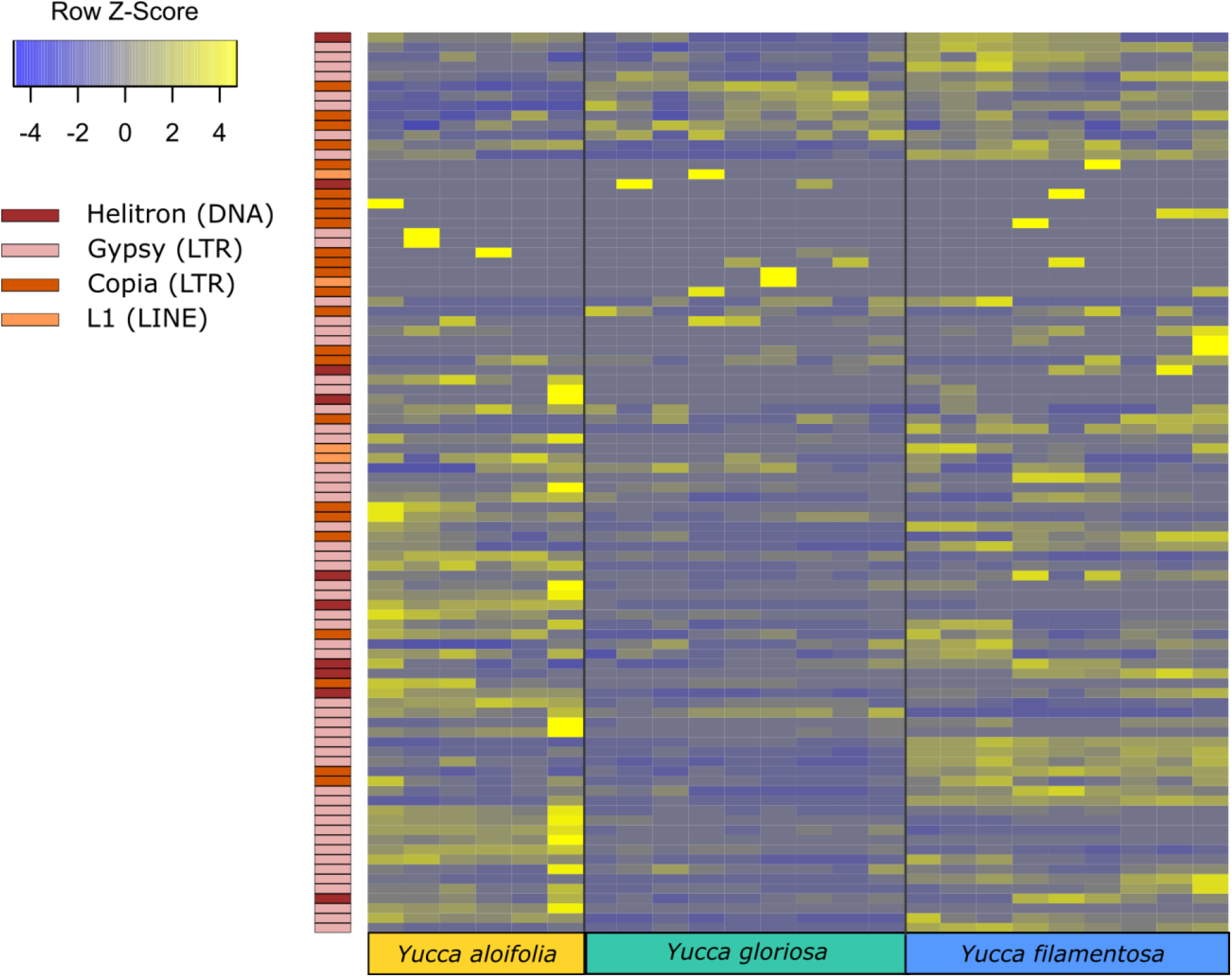
Heatmap of normalized and scaled expression of 92 repeat families that were both present in all three species and had any detectable expression in any library.

**Table 1.**
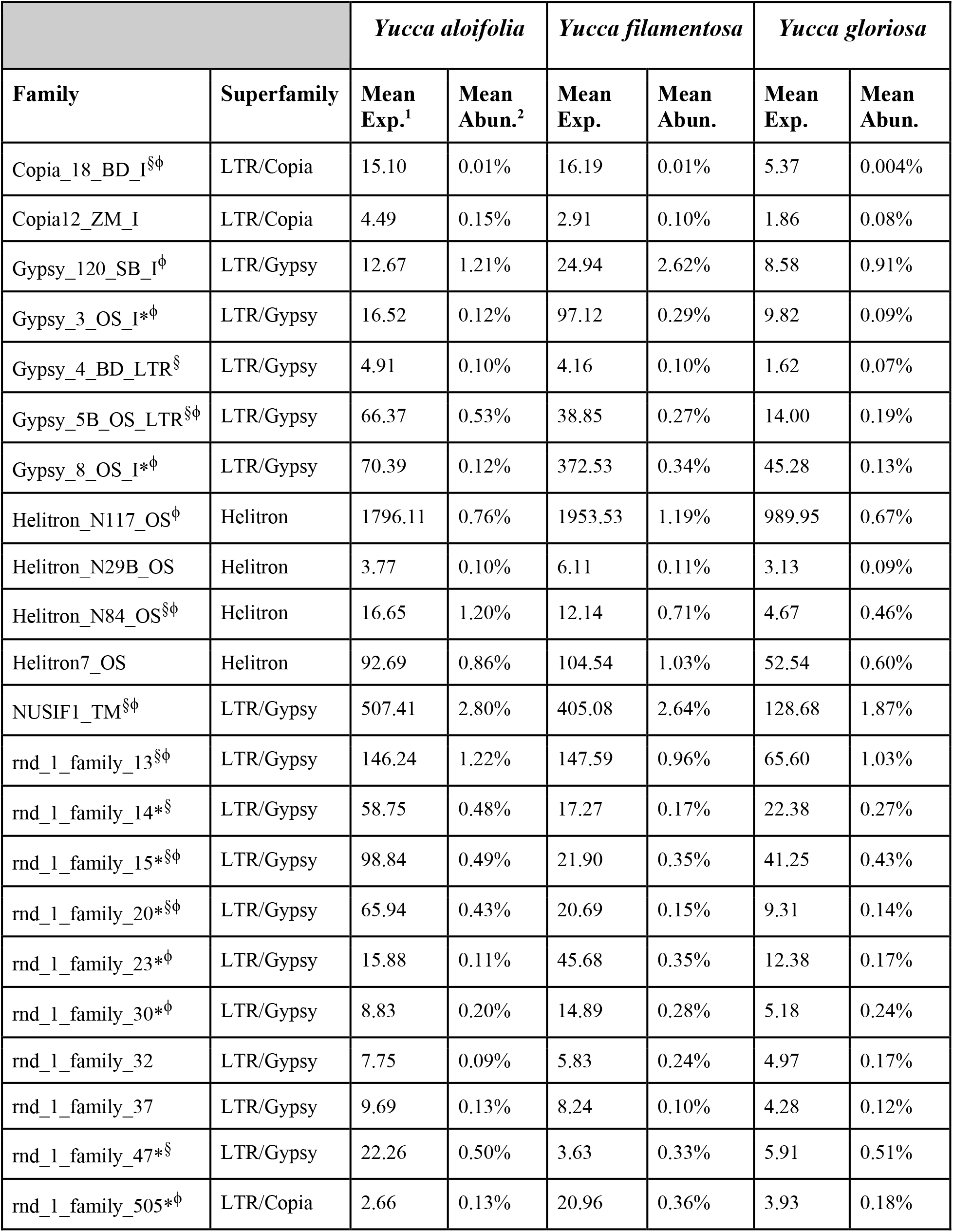
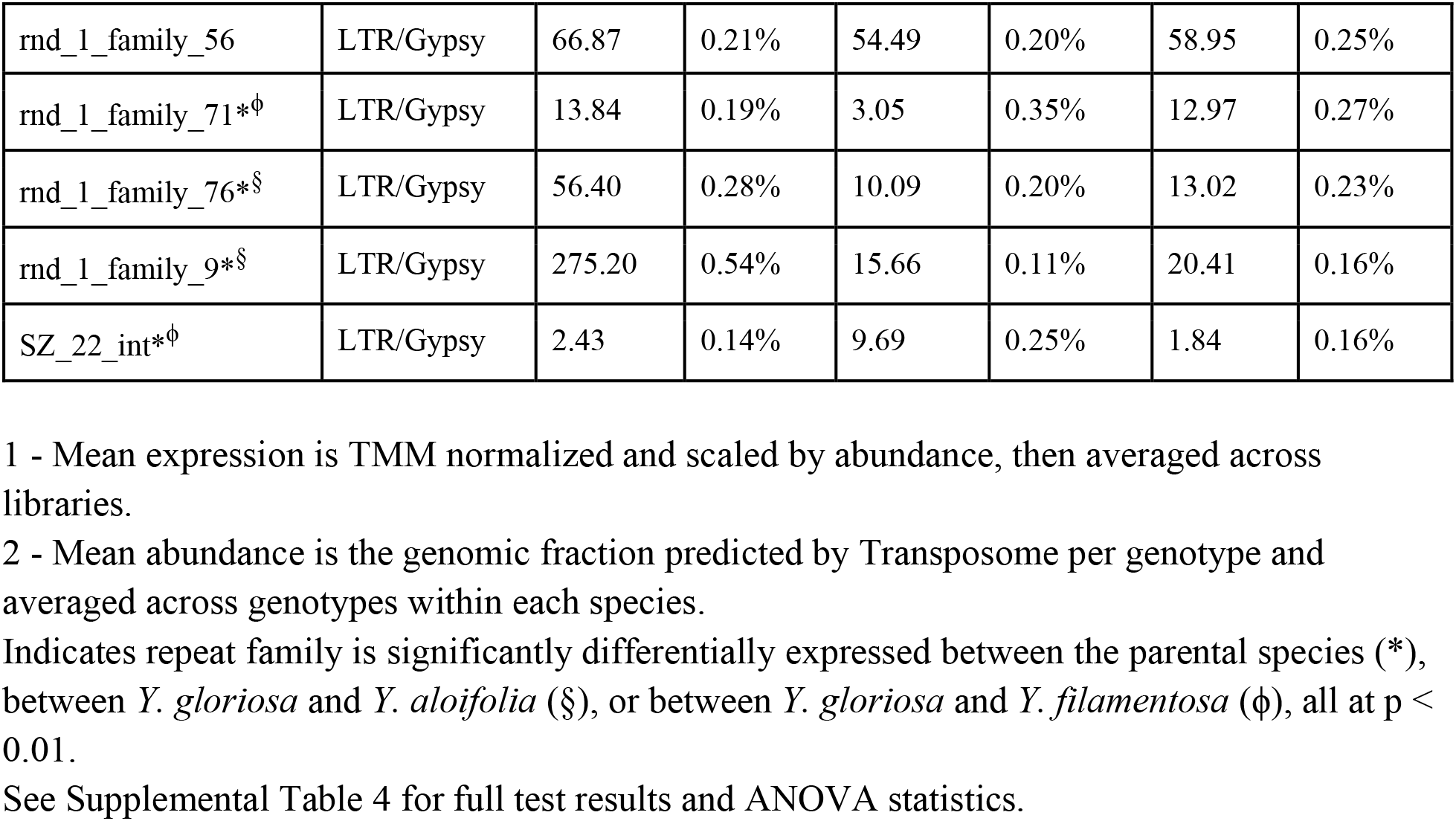
Mean expression and abundance of repeat families in *Y. aloifolia* and *Y. filamentosa* significantly expressed above zero (p<0.01).

|  |  | *Yucca aloifolia* |  | *Yucca filamentosa* |  | *Yucca gloriosa* |  |
| --- | --- | --- | --- | --- | --- | --- | --- |
| Family | Superfamily | Mean Exp.<sup>1</sup> | Mean Abun.<sup>2</sup> | Mean Exp. | Mean Abun. | Mean Exp. | Mean Abun. |
| Copia_18_BD_I<sup>§</sup>φ | LTR/Copia | 15.10 | 0.01% | 16.19 | 0.01% | 5.37 | 0.004% |
| Copia12_ZM_I | LTR/Copia | 4.49 | 0.15% | 2.91 | 0.10% | 1.86 | 0.08% |
| Gypsy_120_SB_I<sup>φ</sup> | LTR/Gypsy | 12.67 | 1.21% | 24.94 | 2.62% | 8.58 | 0.91% |
| Gypsy_3_OS_I*<sup>φ</sup> | LTR/Gypsy | 16.52 | 0.12% | 97.12 | 0.29% | 9.82 | 0.09% |
| Gypsy_4_BD_LTR<sup>§</sup> | LTR/Gypsy | 4.91 | 0.10% | 4.16 | 0.10% | 1.62 | 0.07% |
| Gypsy_5B_OS_LTR<sup>§</sup>φ | LTR/Gypsy | 66.37 | 0.53% | 38.85 | 0.27% | 14.00 | 0.19% |
| Gypsy_8_OS_I*<sup>φ</sup> | LTR/Gypsy | 70.39 | 0.12% | 372.53 | 0.34% | 45.28 | 0.13% |
| Helitron_N117_OS<sup>φ</sup> | Helitron | 1796.11 | 0.76% | 1953.53 | 1.19% | 989.95 | 0.67% |
| Helitron_N29B_OS | Helitron | 3.77 | 0.10% | 6.11 | 0.11% | 3.13 | 0.09% |
| Helitron_N84_OS<sup>§</sup>φ | Helitron | 16.65 | 1.20% | 12.14 | 0.71% | 4.67 | 0.46% |
| Helitron7_OS | Helitron | 92.69 | 0.86% | 104.54 | 1.03% | 52.54 | 0.60% |
| NUSIF1_TM<sup>§</sup>φ | LTR/Gypsy | 507.41 | 2.80% | 405.08 | 2.64% | 128.68 | 1.87% |
| rnd_1_family_13<sup>§</sup>φ | LTR/Gypsy | 146.24 | 1.22% | 147.59 | 0.96% | 65.60 | 1.03% |
| rnd_1_family_14*<sup>§</sup> | LTR/Gypsy | 58.75 | 0.48% | 17.27 | 0.17% | 22.38 | 0.27% |
| rnd_1_family_15*<sup>§</sup>φ | LTR/Gypsy | 98.84 | 0.49% | 21.90 | 0.35% | 41.25 | 0.43% |
| rnd_1_family_20*<sup>§</sup>φ | LTR/Gypsy | 65.94 | 0.43% | 20.69 | 0.15% | 9.31 | 0.14% |
| rnd_1_family_23*<sup>φ</sup> | LTR/Gypsy | 15.88 | 0.11% | 45.68 | 0.35% | 12.38 | 0.17% |
| rnd_1_family_30*<sup>φ</sup> | LTR/Gypsy | 8.83 | 0.20% | 14.89 | 0.28% | 5.18 | 0.24% |
| rnd_1_family_32 | LTR/Gypsy | 7.75 | 0.09% | 5.83 | 0.24% | 4.97 | 0.17% |
| rnd_1_family_37 | LTR/Gypsy | 9.69 | 0.13% | 8.24 | 0.10% | 4.28 | 0.12% |
| rnd_1_family_47*<sup>§</sup> | LTR/Gypsy | 22.26 | 0.50% | 3.63 | 0.33% | 5.91 | 0.51% |
| rnd_1_family_505*<sup>φ</sup> | LTR/Copia | 2.66 | 0.13% | 20.96 | 0.36% | 3.93 | 0.18% |
| rnd_1_family_56 | LTR/Gypsy | 66.87 | 0.21% | 54.49 | 0.20% | 58.95 | 0.25% |
| rnd_1_family_71*<sup>ϕ</sup> | LTR/Gypsy | 13.84 | 0.19% | 3.05 | 0.35% | 12.97 | 0.27% |
| rnd_1_family_76*<sup>§</sup> | LTR/Gypsy | 56.40 | 0.28% | 10.09 | 0.20% | 13.02 | 0.23% |
| rnd_1_family_9*<sup>§</sup> | LTR/Gypsy | 275.20 | 0.54% | 15.66 | 0.11% | 20.41 | 0.16% |
| SZ_22_int*<sup>ϕ</sup> | LTR/Gypsy | 2.43 | 0.14% | 9.69 | 0.25% | 1.84 | 0.16% |

1 - Mean expression is TMM normalized and scaled by abundance, then averaged across libraries.
2 - Mean abundance is the genomic fraction predicted by Transposome per genotype and averaged across genotypes within each species.
Indicates repeat family is significantly differentially expressed between the parental species (\*), between *Y. gloriosa* and *Y. aloifolia* (§), or between *Y. gloriosa* and *Y. filamentosa* (ϕ), all at p < 0.01.
See Supplemental Table 4 for full test results and ANOVA statistics.

In comparing the repeat families that are significantly expressed in *any* of the three species, *Y. gloriosa* showed little transgressive expression patterns; in most of the 178 repeat families that had significant *post hoc* comparisons, *Y. gloriosa* was not statistically different than one of its parental species. There were only three repeat families where expression differed significantly in all three species *(post-hoc* p<0.01) (Supplementary Table 4), and in 5 families, *Y. gloriosa* exhibited an expression level that was significantly different than the pattern shared in the two parental species *(post-hoc* p<0.01) (Fig. 6). In all five cases, *Y. gloriosa* expression was significantly lower than the parental species’ expression, though notably not zero. In general, however, the expression levels of repetitive elements in *Y. gloriosa* were shared with one or both parental species. Nine transposons families showed shared expression in *Y. gloriosa* and *Y. filamentosa* that differed significantly from *Y. aloifolia,* and seven transposons had shared expression between *Y. gloriosa* and *Y. aloifolia* that differed significantly from *Y. filamentosa.* The majority of transposons had shared expression between the two parents, but significantly different expression between *Y. gloriosa* and either *Y. aloifolia* (n=76) or *Y. filamentosa* (n=77). There was a single transposon family where the parental species had significantly different expression from each other and *Y. gloriosa’*s expression was not significantly different than either parent.

**Figure 6.**
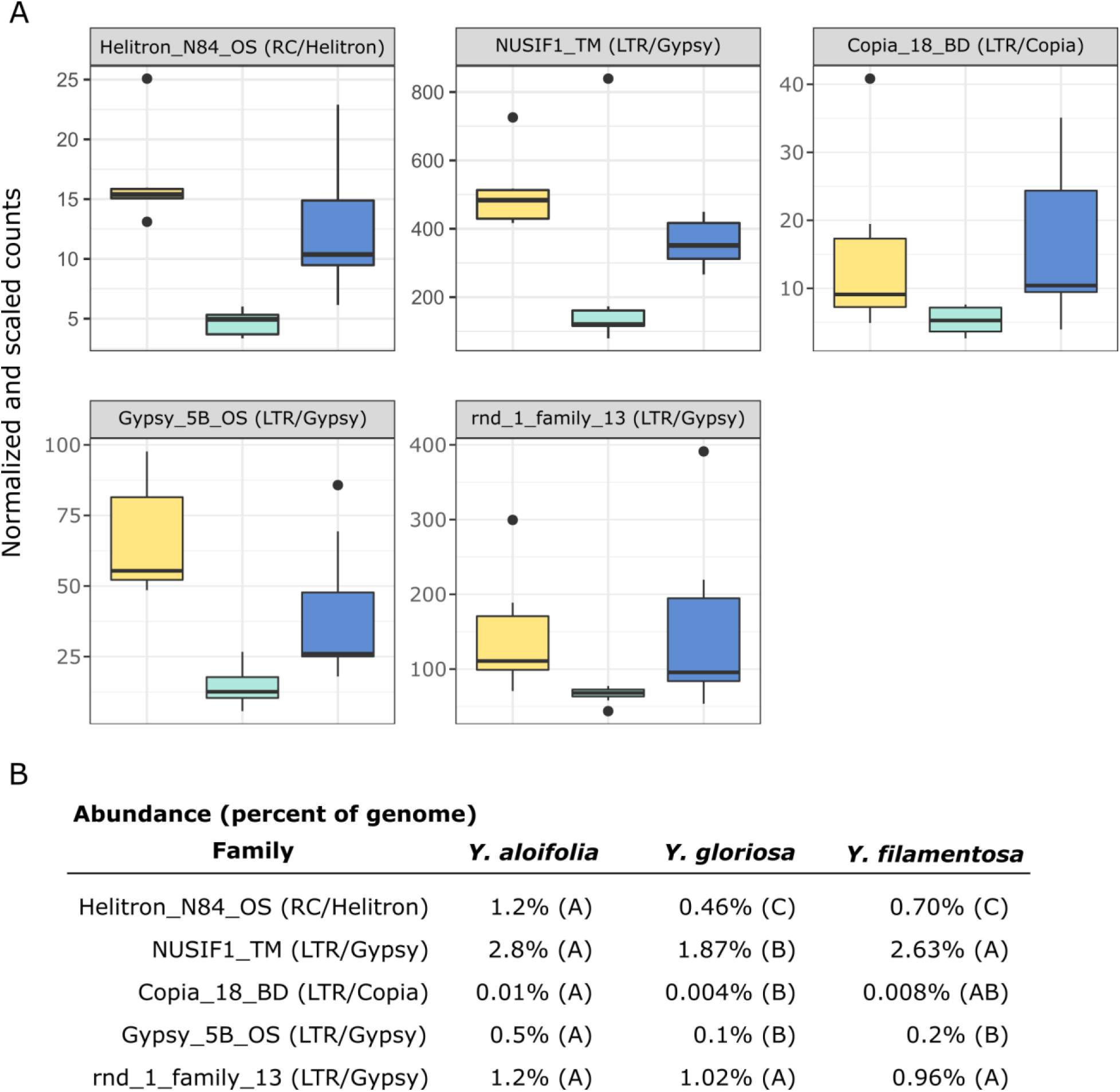
A) Expression plot of the 5 TE families that were significantly differentially expressed between *Y. gloriosa* (teal) and both of its parental species *(Y. aloifolia* = yellow, *Y. filamentosa* = blue). TMM-normalized count data that is further scaled by abundance is plotted. B) Mean percent abundance per species, as estimated by Transposome, and the result of *post hoc* test using emmeans() in R on the results of a negative binomial generalized linear model. Shared letters indicate no significant difference at a p<0.01.

## Discussion

By increasing both the number of *Yucca* genotypes and assessing the whole chloroplast genome we have greatly improved resolution of the history of homoploid hybridization in *Yucca* relative to previous analyses of simple sequence repeats and short fragments of the chloroplast (Rentsch and Leebens-Mack, 2012). Whereas the previous work inferred a single, shared plastid haplotype in *Y. aloifolia* and *Y. gloriosa*, our findings implicate multiple origins of *Y. gloriosa* with both *Y. aloifolia* and *Y. filamentosa* acting as maternal parents. Moreover, analyses of nuclear TE abundances document overall quite similar TE landscapes across the three species, but certain families showed species-specific shifts in abundance. Using mRNA to assess current transposon activity, we find little evidence for ongoing release of transposons in the hybrid genome.

### Reciprocal parentage and multiple origins

Using 15-40x whole genome resequencing data, chloroplast assemblies for 38 individuals of *Yucca* across three species provided robust re-assessment of the history of this hybrid system. The presence of three separate clades containing *Y. gloriosa* (Fig. 2) strongly suggests that not only can *Y. aloifolia* act as the maternal parent in the cross, as previously suggested, but that a reciprocal cross with *Y. filamentosa* as the maternal parent was viable enough to produce at least one extant lineage in *Y. gloriosa*. While *Y. filamentosa* acting as the maternal parent in at least one cross is a parsimonious explanation for the data, the presence of a *Y. filamentosa* chloroplast in *Y. gloriosa* could also be due to a backcrossing event in which a *Y. gloriosa* pollen grain sired a seed on a *Y. filamentosa* individual. Such a backcross is unlikely to have happened recently. Many of the individuals in this study have been phenotyped extensively for photosynthesis related traits (Heyduk et al., 2020), and a recent backcrossed hybrid would be expected to have photosynthetic physiology more similar to *Y. filamentosa* than *Y. aloifolia*, as the parents are strongly divergent in whether they use C3 photosynthesis or CAM, respectively. However, the genotype of *Y. gloriosa* with the *Y. filamentosa* chloroplast haplotype (YG16) has strong signatures of CAM, including nocturnal CO2 uptake as well as acid accumulation, traits which are diagnostic of the CAM phenotype displayed by *Y. aloifolia* (Heyduk et al., 2020). Additionally, the three species are very easy to distinguish in the field by leaf morphology: *Y. filamentosa* has filamentous leaf margins, *Y. aloifolia* has serrated leaf margins, and *Y. gloriosa* has smooth leaf margins. However these observations cannot rule out a more ancient backcrossing event, in which an original *Y. filamentosa* x *Y. gloriosa* cross’ s progeny thereafter crossed only within *Y. gloriosa,* which over time would largely dampen the addition of the *Y. filamenotsa* nuclear genome but the chloroplast haplotype would remain.

The two clades of *Y. gloriosa* individuals that group with *Y. aloifolia* further support that *Y. gloriosa* is derived from multiple hybridization events. However, as with the one instance of a *Y. filamentosa* chloroplast in *Y. gloriosa*, it is difficult to rule out recent backcrossing as the source of this observation (though leaf margins of all *Y. gloriosa* individuals sampled here had smooth margins that are diagnostic of this species in the wild). Additional analysis of reresequencing data will assist in determining the number and timing of putative hybridization events. For example, the length of parental haplotype segments in a hybrid genome is related to the degree of recombination across the hybrid genome; short haplotype blocks would indicate a greater degree of recombination and, therefore, an older hybridization event. On the other hand, longer intact parental haplotype blocks in the hybrid may point to more recent hybridization. Moreover, the length of these haplotype blocks will vary between individuals, and may point to a mixture of both older and younger hybridization events within *Y. gloriosa.*

Previous work on the three *Yucca* species suggested that all *Y. aloifolia* and *Y. gloriosa* individuals shared a single chloroplast haplotype (Rentsch and Leebens-Mack, 2012). Comparisons across the entire chloroplast genome show that four nucleotide differences separated the two clades of *Y. aloifolia* and *Y. gloriosa* individuals. Over 400 genetic changes separate the *Y. filamentosa* and YG16 haplotypes from all *Y. aloifolia* and the remaining *Y. gloriosa* haplotypes. In agreement with the previous work, this study documents low plastid genetic diversity within *Y. aloifolia* and most *Y. gloriosa* samples. *Yucca aloifolia* is introduced into the southeastern United States and likely suffered a bottleneck, resulting in lower overall diversity. The current sample of *Y. gloriosa* individuals identified one individual with a *Y. filamentosa*-derived haplotype. Additionally, this analysis identified seven discrete haplotypes within *Y. filamentosa,* which parallels its greater number of alleles per locus in *Y. filamentosa* suggested by previous work (Rentsch and Leebens-Mack, 2012).

Any attempt at describing the frequency of hybrid formation will be largely affected by the number of individuals in the germplasm collection. The original collection area spanned a large portion of the southeastern United States in order to capture a significant amount of genetic diversity within the genus. Collections of *Y. gloriosa* in particular likely represent many of the extant populations, but the ranges of both *Y. aloifolia* and *Y. filamentosa* are much larger than sampled here. As a result, any interpretation of geographic patterns to the chloroplast phylogeny or haplotype network are hampered by relatively low sampling of the parental genetic diversity. For example, the single *Y. gloriosa* individual found with a *Y. filamentosa* chloroplast (YG16) was collected in South Carolina, while *Y. filamentosa* individuals with the most similar haplotypes were collected in Delaware, North Carolina, and South Carolina. This haplotype grouping is clearly not geographically localized to one portion of the Atlantic coast and could be the result of missing genetic diversity in our analysis. Additionally, the southeastern United States coastline experiences hurricanes and/or tropical storms on nearly an annual basis. Such storms have the potential to both disperse genets as well as eradicate entire populations and could make geographic interpretation of extant diversity difficult.

### Transposable abundance and amplification

Genome resequencing provides a relatively unbiased sampling of the genome, allowing us to estimate the genomic fraction composed of transposable elements. Among sequenced plant genomes, transposable element contribution to genome size ranges from 14% in *Eragrostis tef* to 85% in *Zea mays* (Wendel et al., 2016). While all three *Yucca* species described in this work fall within the described range, the three species varied in the total amount of repetitive DNA with *Y. filamentosa* having significantly less repetitive DNA that *Y. aloifolia* and *Y. gloriosa* (62% vs. 65%/66%). However, variation in abundance of particular repeat superfamilies does suggest superfamily-specific changes between the three species. *Copia* elements, the second most abundant superfamily of repeat in all three species, were more abundant in *Y. gloriosa* relative to both parents, suggesting an amplification of this superfamily post-hybridization. While Class 2 elements represent a relatively small proportion of *Yucca* genomes, *Helitrons* were found more often in *Y. filamentosa* compared to either *Y. aloifolia* or *Y. gloriosa. Helitrons* are capable of generating a tremendous amount of structural novelty, including the ability to capture and redistribute pieces of genes (Yang and Bennetzen, 2009). As genomes become available for these species, it will be possible to analyze the extent to which all types of transposable elements have facilitated structural rearrangements and have affected expression of neighboring genes.

Previous work in various hybrid systems has shown incredible changes to the genomes post-hybridization. In a wallaby x kangaroo cross, reduced methylation of the genome resulted in the proliferation of a novel transposable element that caused significant structural changes to the chromosomes (O’Neill et al., 1998). Interspecific hybrids in *Drosophila* had an increase in transposable element mobilization relative to parental species (Vela et al., 2014). Three independent homoploid hybrids in *Helianthus* all show increased genome size due to expansion of repetitive elements, particularly in *Ty3/gypsy-like* LTR elements (Ungerer et al., 2006, 2009). In *Yucca*, however, there seems to be little indication that transposable elements were released from silencing mechanisms and proliferated in the hybrid *Y. gloriosa*. Instead, *Y. gloriosa* shows similar abundance of transposable elements relative to its progenitor species, though with a notable increase in *Copia* elements in the hybrid (Fig. 4). Extant genotypes of *Y. gloriosa* have little in the way of increased repeat expression (Fig. 6); whether this means no genomic shock initially happened upon hybridization, or that the genome has had sufficient time to stabilize repetitive elements, remains unclear.

Finally, the three *Yucca* species provide an excellent system within which to describe the role of repetitive content on novel phenotypic evolution and adaptation. *Yucca gloriosa* has been studied extensively for its intermediate photosynthetic phenotype (Heyduk et al., 2016, 2019). When well-watered, the majority of carbon fixation happens during the day through the C3 cycle, although low levels of CAM activity are present. When drought stress, *Y. gloriosa* can switch to predominantly CAM photosynthesis, but the degree to which individual genotypes do so varies. The hybrid’s photosynthetic phenotype is novel, in that neither parent displays CAM induction upon drought stress, nor the ability to switch from primarily C3 carbon fixation to primarily CAM. On first glance, negligible differences in repeat content and activity in *Y. gloriosa* relative to its parents suggest that repetitive content is unlikely to underlie the novel photosynthetic phenotype in the hybrid. However, here we only assessed overall abundance and activity in extant individuals; location of repeats in the hybrid relative to the parental species, as well as older repetitive content bursts, still have the potential to create transgressive and novel phenotypes in the hybrid. Repetitive elements can alter gene expression and gene networks by inserting into regulatory regions (Kunarso et al., 2010; Wang et al., 2013), can interfere with alternative splicing (Leprince et al., 2001; Li et al., 2014), and can be a general source of genomic variation and rapid evolution (González et al., 2010; Schrader et al., 2014). Moreover, transposable element activity can increase in response to environmental stressors (Makarevitch et al., 2015) and can play a role in forming stress-induced regulatory networks (Naito et al., 2009). Whether transposable elements are responsible for *Y. gloriosa’*s ability to upregulate CAM photosynthesis under drought stress remains to be tested.

### Conclusions

Since the chloroplast phylogeny and haplotype network imply multiple hybridization events contributing to the origin of *Y. gloriosa,* new hypotheses regarding the repeatability of transposon accumulation can now be tested. For example, since YG16 appears to most likely be derived from a distinct hybridization event relative to other *Y. gloriosa* genotypes, we can assess whether the genomic organization of its transposable elements is vastly different from the major clade of *Y. gloriosa* genotypes grouping with *Y. aloifolia* (Fig. 2). Integrating transposable element abundance and expression with other types of genomic data, including RNA-seq and bisulfite sequencing, may help us understand the potential for insertions to differentially regulate genes. The *Yucca* system is particularly powerful, in that the parental species are strongly divergent in photosynthetic pathway and the hybrid segregates for many of the same traits; this provides a framework in which to understand the role of repeats in regulating these genes in *Y. gloriosa*.

Given the massively expanding availability of whole genome sequence data, hypothesis-driven comparative analyses of genome content and structure are becoming more tractable. In this work, reads that normally would have been filtered out were instead analyzed to address whether a hybrid species had multiple and/or reciprocal origins. Furthermore, these reads helped provide a first glance into the repetitive landscape of 40 genotypes across three related species. While whole genomes will ultimately have the greatest ability to answer many of the questions brought up in this work, the approaches used here are quicker, less expensive, and generate many hypotheses for testing at the genome level in the future.

## Acknowledgements

The authors gratefully acknowledge Amanda L. Cummings for assistance with DNA preparation, the Georgia Advanced Computing Resource Center, and the staff at the University of Georgia greenhouses, in particular Michael Boyd and Gregory Cousins. We thank the DOE Joint Genome Institute and collaborators for pre-publication access to the WGS data from *Yucca* accessions herein and to repeat databases from the genome sequences of *Yucca aloifolia* and *Yucca filamentosa*. This work was supported by a DOE Joint Genome Institute Community Science Project award to K.H. The work conducted by the US DOE Joint Genome Institute is supported by the Office of Science of the US Department of Energy under Contract no. DE-AC02-05CH11231.

## Author Contributions Statement

JG prepared libraries and sequenced samples; SS annotated repetitive content in the parental genomes; EM and KH conducted all analyses and wrote the manuscript; MM optimized plastome assembly and assisted with the manuscript; JLM and JS were integral to overall project planning and management and assisted with the manuscript.

## Conflict of Interest Statement

The authors declare no personal, professional, or financial conflicts of interest.

## Notes

### Competing Interest Statement

The authors have declared no competing interest.

