## Supplementary Figure 1 for "Hybridization history and repetitive element content in the genome of a homoploid hybrid, *Yucca gloriosa* (Asparagaceae)"

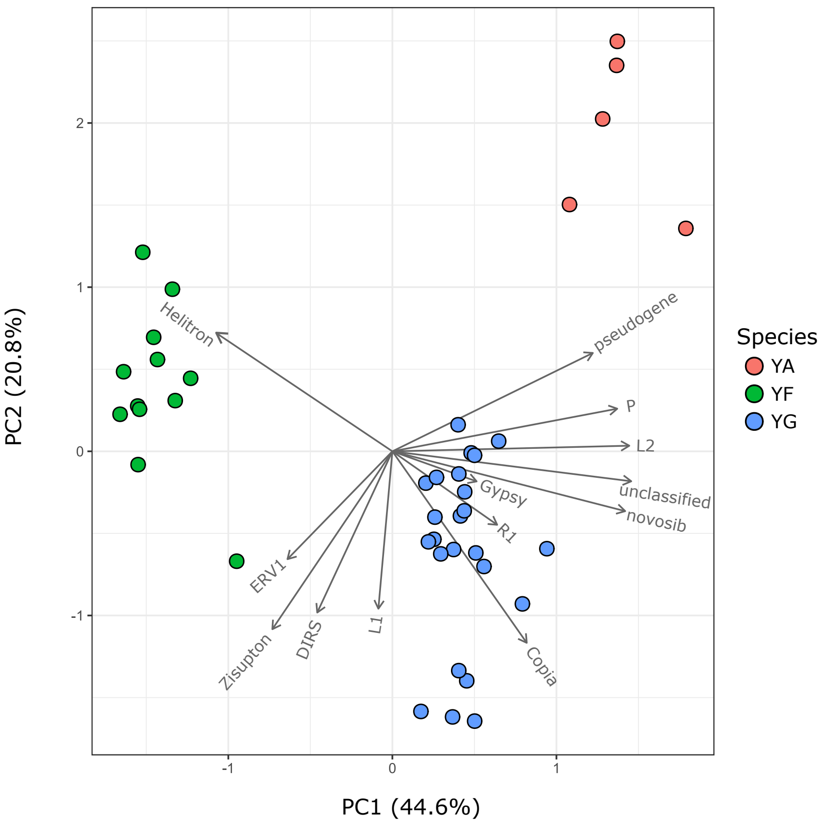


**Supplemental Figure 1** - Principal Coordinates Analysis of repeat subfamily abundance in genotypes of three *Yucca* species.
